# Maternal Ezh1/2 deficiency in oocyte delays H3K27me2/3 restoration and impairs epiblast development responsible for embryonic sub-lethality in mouse

**DOI:** 10.1101/2021.10.28.466222

**Authors:** Yinan Zhao, Dan Zhang, Mengying Liu, Yingpu Tian, Jinhua Lu, Shaorong Gao, Haibin Wang, Zhongxian Lu

## Abstract

Mammalian embryonic development is a complex process regulated by various epigenetic modifications. Recently, maternal histone H3 methylations were found to be inherited and reprogrammed in early embryos to regulate embryonic development. The enhancer of zest homolog 1 and 2 (Ezh1 and Ezh2) belong to the core components of Polycomb repressive complex 2 (PRC2) and are the histone methyltransferase of histone 3 lysine 27 (H3K27). How maternal Ezh1 and Ezh2 function on H3K27 methylation in *in vivo* preimplantation embryos and embryonic development are not clear. Here, we deleted *Ezh1* or/and *Ezh2* in growing oocytes using gene knockout mouse models, and found that H3K27me3 in oocytes was disappeared by loss of Ezh2 alone while H3K27me2 was absent upon deletion of both Ezh1 and Ezh2. The effects of *Ezh1/2* were inherited in maternal knockout zygotes and early embryos, in which restoration of H3K27me3 was delayed until late blastocyte by loss of Ezh2 alone and H3K27me2 was reestablished until morulae by deletion of Ezh1 and Ezh2. However, the ablation of both *Ezh1* and *Ezh2*, but not single *Ezh1* or *Ezh2*, led to significantly decreased litter size due to growth retardation during post-implantation. Furthermore, maternal *Ezh1/2* deficiency caused compromised H3K27me3 and pluripotent epiblast cells in late blastocyst, followed by defective development of epiblast. These results demonstrate that in oocytes, *Ezh2* is indispensable for H3K27me3 while *Ezh1* complements *Ezh2* in H3K27me2. Also, maternal Ezh1/2-H3K27 methylation is inherited in descendant embryos and has a critical effect on fetus and placenta development. Thus, this work sheds light on maternal epigenetic modifications during embryonic development.

## Introduction

Mammalian embryonic development begins with the formation of a zygote and ends when a fully developed fetus is delivered. This colorful process is accompanied with a variety of epigenetic modifications (Saitou et al., 2012). Histone modifications are found to play key roles in epigenetic regulation (Canovas and Ross, 2016). The methylations of lysine 4, 9, 27 on histone H3 are transcriptional modulators that regulate specific sets of genes involved in embryogenesis, embryonic stem cell (ESC) pluripotency and differentiation, maintenance of tissue stem cell, establishment and maintenance of genomic imprinting, as well as tumorigenesis (Jambhekar et al., 2019; Michalak et al., 2019; Yu et al., 2019). Recently, dynamic patterns of H3K4 trimethylation (H3K4me3) and H3K27 trimethylation (H3K27me3) which play critical roles in maternal X-chromosome inactivation (XCI), zygotic genome activation (ZGA) and developmental genes, are illustrated in mouse oocytes and preimplantation embryos (Inoue et al., 2018; Liu et al., 2016; Zhang et al., 2016). Histone modifications are germ line inherited or reestablished during early embryogenesis (Liu et al., 2016; Zhang et al., 2016; Zheng et al., 2016). However, the regulation and underlying mechanism of maternal histone modifications in embryonic development need to be fully explored.

PRC2 is a histone methyltransferase (HMT) complex to produce cell type specific H3K27 mono- (H3K27me1), di- (H3K27me2), and tri-methylation (H3K27me3) patterns (Bracken et al., 2006; Ferrari et al., 2014). H3K27me3 is a well-known repressive histone modification ultimately resulting in transcriptional silence (Cao et al., 2002). In mammals, the core subunits of PRC2 complex include enhancer of zeste homolog 1 or 2 (Ezh1 or Ezh2), embryonic ectoderm development (Eed), suppressor of zeste 12 (Suz12) and retinoblastoma binding protein 4/7 (Rbbp4/7) (Yan et al., 2019). Ezh1 and Ezh2 are the catalytic subunits of the PRC2 complex (Margueron and Reinberg, 2011). In *Drosophila*, deletion of Enhancer of zeste [E(z)] in oogenesis causes embryonic lethality although H3K27me3 is reestablished in late zygotes (Zenk et al., 2017), indicating H3K27me3 requires maternal PRC2 during early embryogenesis. But, the effects of maternal PRC2 on mammalian embryonic development remain unclear and inconsistent. In mice, oocyte-specific deletion of *Ezh2* by *Zp3*-*Cre* shows no change in litter size but growth retardation in progeny (Erhardt, 2003). Maternal *Eed* knockout by *Gdf9*-*Cre* shows loss of H3K27me3 imprinting and growth arrest in post-implantation (Inoue et al., 2018). Loss of *Eed* from growing oocytes by *Zp3*-*Cre* results in a significant overgrowth phenotype that persisted into adult life (Prokopuk et al., 2018). These results indicate that *Ezh2* and *Eed* may have different functions in post-embryonic development (Prokopuk et al., 2018). Collectively, it seems that maternal PRC2 core components have a long-term effect on descendant development. But the exact details need to be discovered. Recently, a report shows that Ezh2 is required for the establishment of H3K27me3 in mouse zygotes, Ezh1 could partially safeguard the role of Ezh2 (Meng et al., 2020), indicating that *Ezh1* might compensate for *Ezh2* in specific developmental systems (Ezhkova et al., 2011; Mochizuki-Kashio et al., 2015). However, how maternal Ezh1 and Ezh2 comprehensively mediate H3K27 methylation in *in vivo* preimplantation embryos is unknown.

To this end, we use maternal *Ezh1/2* knockout mice to gain insights into the dynamic patterns of H3K27 methylation during preimplantation and reveal their regulation on the embryonic development. The results demonstrated that maternal *Ezh1/2* are required for the establishment of H3K27me2/3 in *in vivo* preimplantation embryos and play critical roles in embryonic development in mouse.

## Results

### Recovery of H3K27me3 and H3K27me2 are delayed in early maternal knockout embryos

To reveal the maternal PRC2 mediated histone modifications and its long-term effect on embryonic development, we introduced an Ezh1 null mouse line and Gdf9-Cre transgenic mice to construct transgenic mouse models with *Ezh1* and *Ezh2* deletion in growing oocytes. Here, we established two pairs of mouse lines, one pair were sCtrl (single control) and sKO (single knockout of Ezh2 in oocyte with Gdf9-Cre) **(sFig. 1A)**, and the other were dCtrl (double control, that is *Ezh1* knocked out) and dKO (double knockout, that is both *Ezh1* and *Ezh2* knocked out in oocytes) **(sFig. 2A)**. In sKO mice, the mRNA and protein of Ezh2 in oocytes was undetectable **(sFig. 1B and 1C)**, whereas the expression of other PRC2 core components, Ezh1, Eed, and Suz12 was not affected **(sFig. 1B)**. In dKO mice, *Ezh1* and *Ezh2* genes were deleted while the mRNA level of *Eed* and *Suz12* was not affected **(sFig. 2B and 2C)**. These observations showed that Ezh1 and Ezh2 in oocytes were successfully deleted in these mice.

PRC2 complex is responsible for methylations on H3K27 through its enzymatic subunits Ezh1 and Ezh2 (Schuettengruber and Cavalli, 2009; Simon and Kingston, 2009). Then, we checked H3K27me2/3 modifications in oocytes by immunohistochemistry (IHC) and Immunofluorescence (IF). H3K27me3 was absent in oocytes from secondary follicles and MII oocytes from both sKO **(sFig. 1D and 1E)** and dKO mice **(sFig. 2D-2F)**. But, H3K27me2 was disappeared only in MII oocytes of dKO mice, neither in sKO nor in dCtrl mice **(sFig. 2F)**. These results suggest that Ezh2 is indispensable for H3K27me3 while Ezh1 and Ezh2 are complementary on H3K27me2 modification in oocytes.

Then, we examined H3K27 methylation patterns at different stage of early embryos. Embryos from sCtrl, sKO, dCtrl and dKO females mated with wild type males were referred to as sF/+, sKO/+, dF/+ and dKO/+ respectively hereafter. H3K27me3 staining was strongly in female pronucleus (PN) but relatively weak in paternal PN in sF/+ late zygotes **(Fig. 1A)**. From late 2-cell stage to early blastocysts, H3K27me3 was clearly detected in sF/+ embryos, while H3K27me3 staining was not restored until late blastocysts in sKO/+ groups **(Fig. 1A)**, suggesting the modification of H3K27me3 in preimplantation embryos is dependent on maternal Ezh2. Enhanced H3K27me3 staining in inner cell mass (ICM) appeared at sKO/+ late blastocyst stage, along with dot staining in trophectoderm (TE) in some embryos, which was comparable to sF/+ embryos **(Fig. 1A)**. In addition, H3K27me3 staining was still faint in late blastocysts of dKO/+ embryos **(Fig. 1B**, right images), indicating that maternal Ezh1 assists Ezh2 in H3K27me3 modification in early embryos.

**Figure 1.**
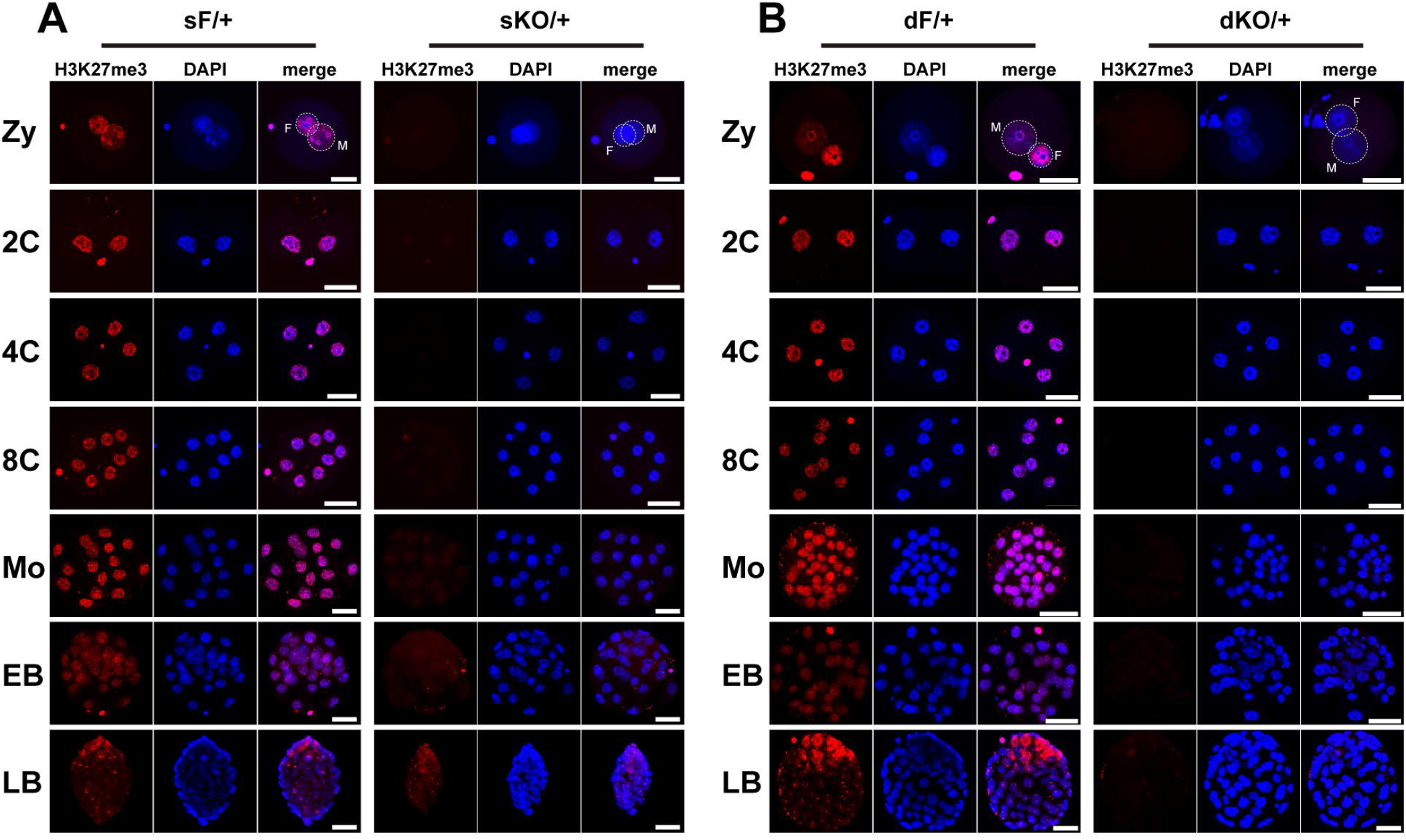
Delayed recovery of H3K27me3 modification in preimplantation stage of sKO/+ and dKO/+ embryos. **(A)** H3K27me3 patterns in sF/+ and sKO/+ embryos. In late zygotes, H3K27me3 was absent in both male and female pronucleus (PN) of sKO/+. H3K27me3 was not detectable until late blastocysts in sKO/+ embryos. The number of zygote examined: sF/+, n=22; sKO/+, n=21. 2-cell embryo: sF/+, n=21; sKO/+, n=25. 4-cell embryo: sF/+, n=32; sKO/+, n=12. 8-cell embryo: sF/+, n=12; sKO/+, n=16. Morula: sF/+, n=19; sKO/+, n=17. Early blastocyst: sF/+, n=18; sKO/+, n=14. Late blastocyst: sF/+, n=5; sKO/+, n=11. Scale bars, 50 μm. **(B)** H3K27me3 patterns in dF/+ and dKO/+ embryos. H3K27me3 was not detectable even in dKO/+ late blastocysts. The number of zygote examined: dF/+, n=19; dKO/+, n=14. 2-cell embryo: dF/+, n=25; dKO/+, n=18. 4-cell embryo: dF/+, n=20; dKO/+, n=17. 8-cell embryo: dF/+, n=19; dKO/+, n=14. Morula: dF/+, n=35; dKO/+, n=38. Early blastocyst: dF/+, n=13; dKO/+, n=11. Late blastocyst: dF/+, n=4; dKO/+, n=12. Scale bars, 50 μm. Zy: zygote (PN4-PN5); 2C: 2-cell embryo; 4C: 4-cell embryo; 8C: 8-cell embryo; Mo: morula; EB: early blastocyst; LB: late blastocyst; M: male pronucleus; F: female pronucleus.

During normal embryonic development, H3K27me2 has a similar expression pattern to that of H3K27me3. In sKO/+ embryos, H3K27me2 in female PN was very weak in contrast to that of sF/+, and was almost undetectable in paternal PN **(Fig. 2A**, right images). H3K27me2 was detected at 2-cell stage in sKO/+ embryos and comparable to sF/+ embryos from 4-cell to late blastocyst stage **(Fig. 2A)**. However, H3K27me2 staining is absent from zygotes to 8-cell stage in dKO/+ embryos, faint from morulae to early blastocysts, and fully recovered until late blastocysts **(Fig. 2B**, right images), indicating maternal Ezh1 in conjunction with Ezh2 plays an important role in H3K27me2 modification. Intriguingly, the dot staining of H3K27me2 was strongly increased in dKO/+ embryos and some cells had two dot staining at morula stage **(sFig. 3A and 3B)**, unveiling that H3K27me2 may be contributed to X-chromosome inactivation (XCI).

**Figure 2.**
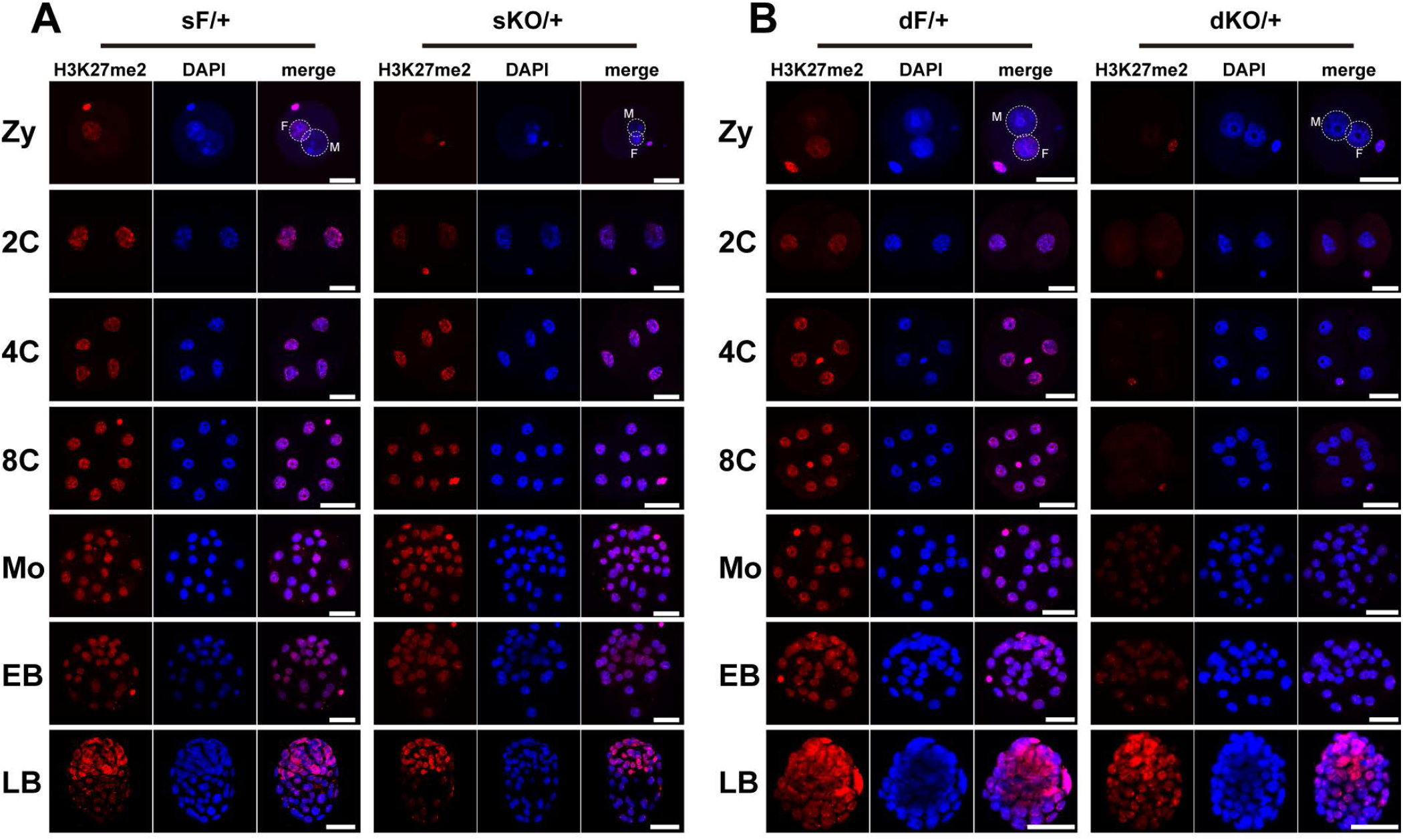
Delayed recovery of H3K27me2 modification in preimplantation stage of sKO/+ and dKO/+ embryos. **(A)** H3K27me2 patterns in sF/+ and sKO/+ embryos. In sKO/+ late zygotes, H3K27me2 was undetectable in male PN and faint in female PN. H3K27me2 was recovered completely in 4-cell embryos in sKO/+. The number of zygote examined: sF/+, n=21; sKO/+, n=20. 2-cell embryo: sF/+, n=40; sKO/+, n=29. 4-cell embryo: sF/+, n=25; sKO/+, n=16. 8-cell embryo: sF/+, n=36; sKO/+, n=44. Morula: sF/+, n=20; sKO/+, n=16. Early blastocyst: sF/+, n=12; sKO/+, n=5. Late blastocyst: sF/+, n=13; sKO/+, n=6. Scale bars, 50 μm. **(B)** H3K27me2 patterns in dF/+ and dKO/+ embryos. H3K27me2 in dKO/+ embryos was not detectable until the morula stage. Note dot signals in morulae and blastocysts. The number of zygote examined: dF/+, n=17; dKO/+, n=10. 2-cell embryo: dF/+, n=32; dKO/+, n=38. 4-cell embryo: dF/+, n=22; dKO/+, n=23. 8-cell embryo: dF/+, n=11; dKO/+, n=14. Morula: dF/+, n=24; dKO/+, n=25. Early blastocyst: dF/+, n=12; dKO/+, n=5. Late blastocyst: dF/+, n=4; dKO/+, n=15. Scale bars, 50 μm. Zy: zygote (PN4-PN5); 2C: 2-cell embryo; 4C: 4-cell embryo; 8C: 8-cell embryo; Mo: morula; EB: early blastocyst; LB: late blastocyst; M: male pronucleus; F: female pronucleus.

Together, these observations revealed the dynamic patterns of maternal H3K27 methylation during preimplantation and demonstrate that H3K27 methylation activated by maternal PRC2 is dominant in preimplantation embryos. Maternal Ezh2 is required for H3K27me3 modification and maternal Ezh1 plays assistant roles in H3K27me2/3 modification in oocyte and early embryo.

### Double knockout of *Ezh1* and *Ezh2* in oocytes results in female subfertility

Then, we investigated the reproductive performance of the KO mice. Females were mated with wild type (WT) males. The cumulative number of pups, number of pups per litter and the number of litters per mouse were analyzed. The results showed sKO females had normal fertility **(Fig. 3A-3C)**. While cumulative number of pups produced from dKO females was obviously lower than that of dCtrl females **(Fig. 3D)**, the average litter size from dKO females (2.69 ± 1.35) was significantly smaller compared to dCtrl females (7.10 ± 2.13) **(Fig. 3E)**. The number of litters per mouse in dKO females was also greatly decreased **(Fig. 3F)**. These data discovered that loss of both Ezh1 and Ezh2 in oocytes impaired female fertility in mouse, indicating that maternal PRC2 has a long-term effect on embryonic development and maternal Ezh2 combined with Ezh1 governs the descendant development.

**Figure 3.**
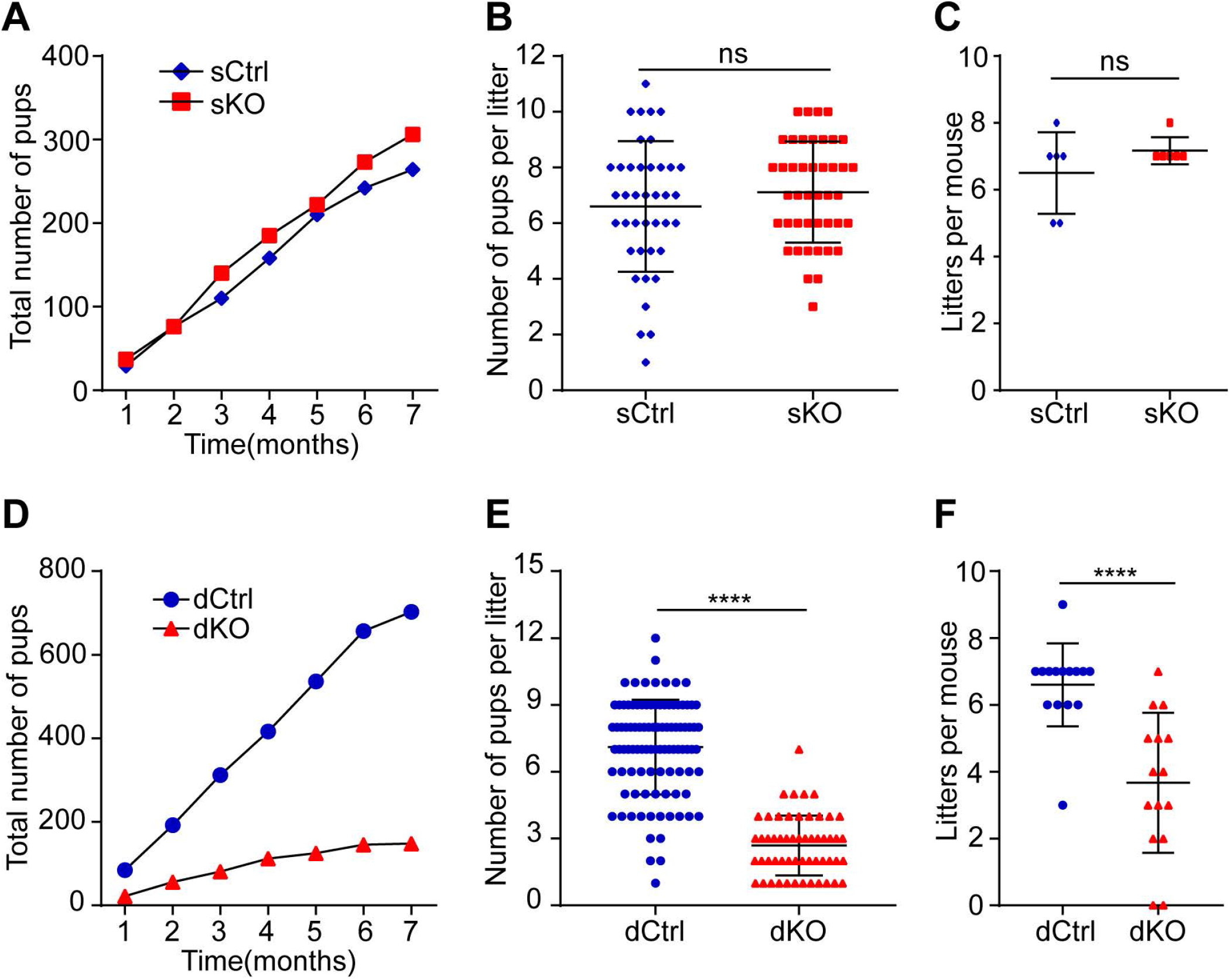
Double knockout of *Ezh1* and *Ezh2* in oocytes leads to female subfertility. **(A-C)** Fertility studies of sCtrl and sKO mice. Number of females: sCtrl, n=6; sKO, n=6. **(A)** The cumulative number of pups. **(B)** The average number of pups per litter. Data are presented as the mean ± SD. Mann Whitney test: ns, not significant. **(C)** The average number of litters per mouse. Data are presented as the mean ± SD. Mann Whitney test: ns, not significant. **(D-F)** Fertility studies of dCtrl and dKO mice. Number of females: dCtrl, n=15; dKO, n=15. **(D)** The cumulative number of pups. **(E)** The average number of pups per litter. Data are presented as the mean ± SD. Mann Whitney test: \*\*\*\**P* < 0.0001. **(F)** The average number of litters per mouse. Data are presented as the mean ± SD. Mann Whitney test: \*\*\*\**P* < 0.0001.

### Maternal Ezh1/2 knockout embryos develop abnormally after implantation

Since histone modifications are reprogrammed widely during gametogenesis (Matsui and Mochizuki, 2014; Zheng et al., 2016), ovary development and ovulation were firstly examined to reveal the reason responsible for subfertility. In dKO females, ovarian morphology was not affected and follicles developed normally **(sFig. 3A)**. The average number of ovulated oocytes in dKO groups was also similar to that of dCtrl mice **(sFig. 3B and 3C)**. Then, there was no difference in the number of implantation sites on day 5 of pregnancy between dKO and dCtrl females **(sFig. 3D and 3E)**, indicating that preimplantation development and embryo implantation are normal. These results suggest that decreased fertility is not due to oogenesis and implantation.

Then, we focused on embryos in post-implantation and isolated embryos at E10.5, when the mouse embryo is under organogenesis (Kojima et al., 2014) and the basic structure of mouse placenta is formed (Rossant and Cross, 2001). Morphologically, the dKO/+ fetus and placenta appeared severely retarded at E10.5 **(Fig. 4A)**. Partial embryos (26.83%) had been absorbed and failed to isolate **(Fig. 4B)**. In addition, the decidua weight of dKO females was greatly reduced compared to that of dCtrl females **(Fig. 4C)**, indicating growth restriction. HE staining showed that phenotypes of dKO/+ embryos at E10.5 could be identified into three categories. Type I had a placenta containing labyrinth (Lab) layer, spongiotrophoblast (Sp) layer (always smaller) and trophoblast giant cells (TGC) **(Fig. 4D**, left second images); type II had fetus while had no Lab and Sp **(Fig. 4D**, right second images); and embryos from type III were in absorption or have been absorbed **(Fig. 4D**, right first images).

**Figure 4.**
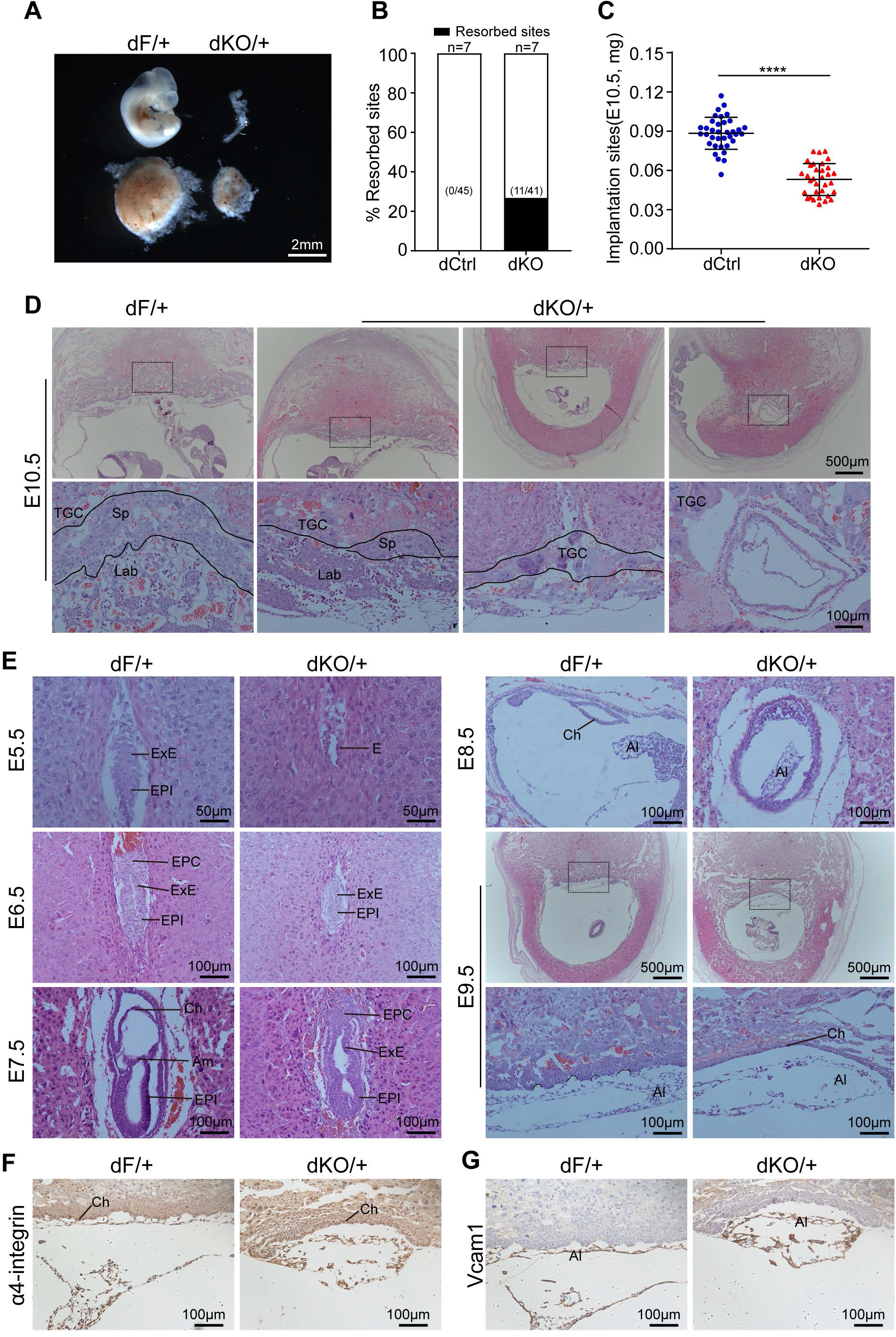
Double knockout of *Ezh1* and *Ezh2* in oocytes causes growth retardation during post-implantation. **(A)** Representative images of E10.5 fetuses (top) and placentae (bottom). **(B)** Ratio of resorbed sites at E10.5 in dCtrl and dKO females. Numbers within parentheses indicated the number of resorbed sites over total number of implantation sites. Number of females examined was shown on top of the bars. **(C)** The weight of implantation sites at E10.5. Number of implantation sites: dCtrl, n=36; dKO, n=33. Results are presented as mean ± SD. Unpaired *t* test: \*\*\*\**P* < 0.0001. **(D)** Representative images of placentae at E10.5 by HE staining. Black rectangles indicated the enlarged regions shown in the below panels. **(E)** Representative images of uterine decidua sections from E5.5 to E9.5 stained with HE. Black rectangles indicated the enlarged regions shown in the below panels. Lines at E9.5 in left bottom indicated progression of labyrinth branching and the image in right bottom showed that branching was failed. **(F)** The expression of Integrin α4 in chorion at E9.5 by IHC staining. **(G)** Vcam1 expression in allantois at E9.5 by IHC staining. TGC, trophoblast giant cells; Sp, spongiotrophoblast; Lab, labyrinth; ExE, extraembryonic ectoderm; EPI, epiblast; E, embryo; EPC, ectoplacental cone; Ch, chorion; Al, allantois.

Furthermore, growth retardation of embryos from E5.5-E7.5 was observed in decidua sections **(Fig. 4E**, E5.5-E7.5). Indeed, paraffin section and dissection under stereoscope showed that embryos diminished from E5.5. Moreover, failure of chorioallantoic attachment and branching was observed in dKO/+ embryos at E9.5, in line with placental defects at E10.5 **(Fig. 4E)**. This demonstrates that growth arrest during post-implantation is responsible for subfertility. It has been known that chorioallantoic attachment during placental development is dependent on Vcam1, a cell-adhesion molecule expressed on the tip of the allantois, and its ligand integrin α4, expressing on the basal surface of the chorion (Rossant and Cross, 2001). However, IHC analysis showed that Vcam1 and integrin α4 expressed normally in dKO/+ embryos **(Fig. 4F and 4G)**, indicating that the interactions between Vcam1 and integrin α4 alone is not sufficient for a successful union of chorion and allantois (Walentin et al., 2016) and other factors may account for failed chorioallantoic attachment observed in dKO/+ embryos.

Surprisingly, morphological and histological analyses revealed that the placentas of dKO/+ embryos were enlarged in late stage of pregnancy **(Fig. 5A and 5B)**, which was shown by the overgrowth of spongiotrophoblast and increased placenta weight at E17.5 **(Fig. 5C and 5F)**. However, fetuses were smaller than dF/+ groups **(Fig. 5A)**, which were exhibited by the decreased weight and length of fetus **(Fig. 5D and 5E)**.

**Figure 5.**
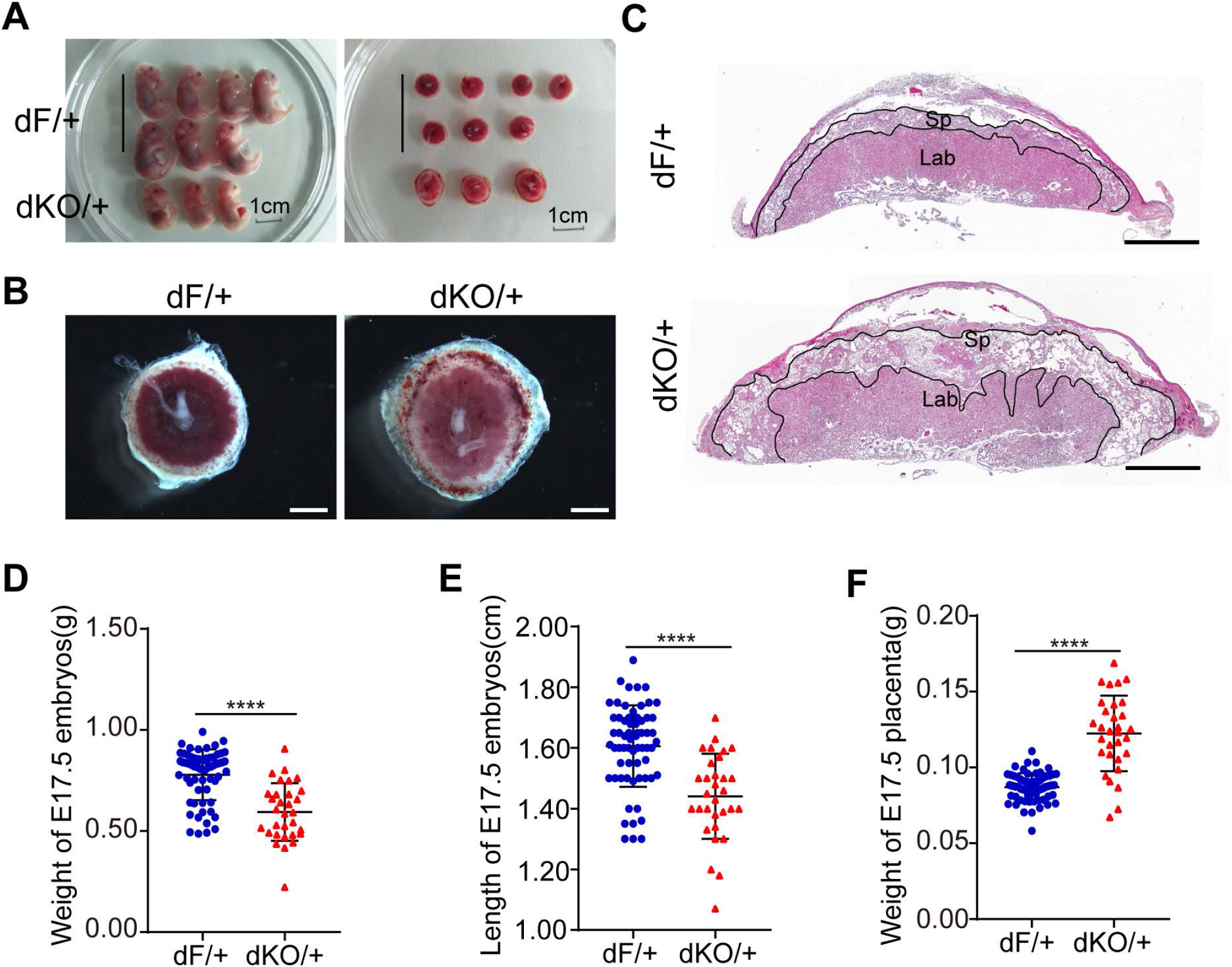
The enlarged placentae in late stage of dKO/+ embryos. **(A)** Representative images of E17.5 fetuses and placentae. The top seven showed the dF/+ embryos and the bottom three showed the dKO/+ embryos. **(B)** Representative images of E17.5 pacentae under the stereoscope. Scale bars, 1 mm. **(C)** Morphology of E17.5 placentae by HE staining. Sp, spongiotrophoblast; Lab, labyrinth. Scale bars, 1 mm. **(D)** The weight of fetus at E17.5. Results are presented as mean ± SD. Mann Whitney test: \*\*\*\**P* < 0.0001. **(E)** The length of fetus at E17.5. Results are presented as mean ± SD. Mann Whitney test: \*\*\*\**P* < 0.0001. **(F)** The weight of placenta at E17.5. Results are presented as mean ± SD. Unpaired *t* test: \*\*\*\**P* < 0.0001.

In summary, the defective development of embryos fell into three distinct categories. The first was that embryos ceased development shortly after implantation and could not exceed gastrulation; the second was that embryos with failure of placentation were destined to death; the third was that embryos could overcome gastrulation and placentation, but accompanied by giant placenta and smaller fetus.

To determine whether maternal uterine environment contributes to growth retardation, embryo transfer experiments were pursued. dF/+ and dKO/+ blastocysts were transferred to WT pseudo-pregnant recipients separately. dKO/+ groups showed considerably reduced number of term pups (**Supplemental Table 1**). This result is consistent with subfertility of dKO females. About 18% of dKO/+ blastocysts developed to term pups, much lower than that of dF/+ blastocysts (about 48%) (**Supplemental Table 1**). This indicates that dKO/+ embryo *per se* rather than maternal uterine environment that contributes to abnormal embryonic development.

Above all, these results suggest that maternal Ezh1/2 deficiency in oocytes disturbs the embryonic development and loss of maternal Ezh1/2 has a long-lasting consequence on embryogenesis.

### EPI cells are abnormal in dKO/+ embryos during peri-implantation

To trace the onset and progression of developmental abnormalities, the development of dKO/+ embryos were examined during peri-implantation in mice. Normally, epiblast (EPI) cells differentiated from the inner cell mass (ICM) proliferate rapidly after implantation, and then the embryos form pro-amniotic cavity at E5.5, a prerequisite for gastrulation (Mole et al., 2020; Tam and Loebel, 2007). However, no discernible epiblast was observed in dKO/+ embryos at E4.75 **(Fig. 6A)**. Successive sections showed that dKO/+ embryos had few Oct4+ cells at E4.75 **(Fig. 6B)**. At E5.5, dKO/+ embryos had no pro-amniotic cavity while dF/+ embryos showed an obvious pro-amniotic cavity **(Fig. 6C)**. Our careful morphological examinations at different embryonic stages reveal that growth arrest appears during peri-implantation and the EPI/ICM development is defective.

**Figure 6.**
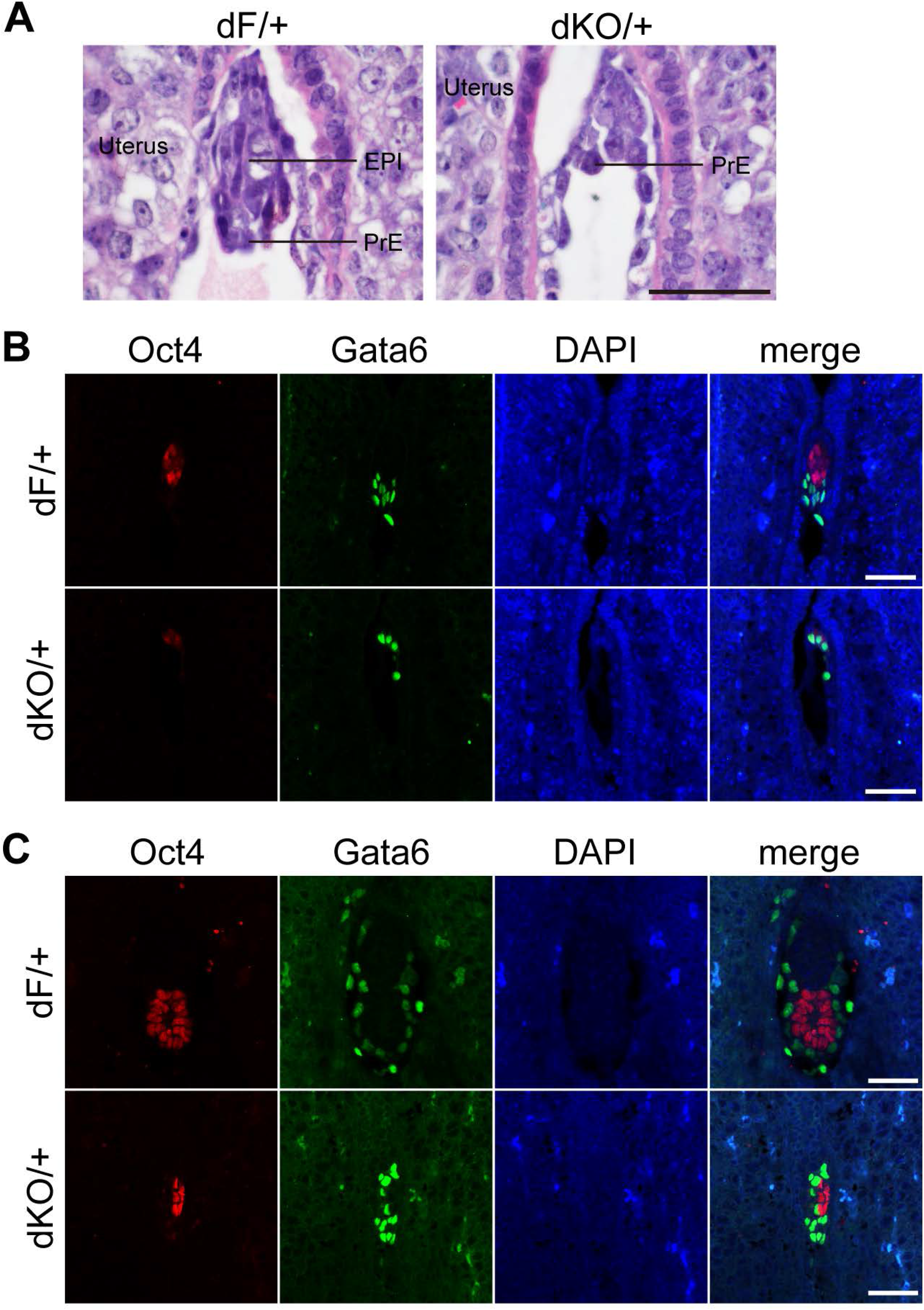
The defective development of epiblast in post-implantation dKO/+ embryos. **(A)** Uterine decidua sections by HE staining at E4.75. Scale bars, 50 μm. EPI: epiblast; PrE: primitive endoderm. **(B)** The expression of Oct4 and Gata6 in embryos at E4.75 by IF staining. Scale bars, 50 μm. **(C)** The expression of Oct4 and Gata6 in embryos at E5.5 by IF staining. Scale bars, 50 μm. Epiblast (EPI) cells were marked by Oct4 (red). Primitive endoderm (PrE) cells, visceral endoderm (VE) cells and parietal endoderm (PE) cells were marked by Gata6 (green).

We next sought to determine how the EPI/ICM cells are impaired. In mouse development, the first cell lineage decision occurs in preimplantation (around at E3.0~3.5) and generates two cell populations: TE cells expressing the *Cdx2* genes and ICM cells expressing the *Oct4* gene (Chazaud and Yamanaka, 2016). We examined the first cell fate decision by immunostaining with Oct4 and Cdx2 at E3.75. There was no different in the allocation of TE and ICM cells between dF/+ and dKO/+ embryos **(Fig. S5A)**. Neither the number of ICM and TE cells nor their ratios were significantly changed, although both-expressed cells were increased in dKO/+ embryos **(Fig. S5B and 5C)**. It’s shown that the first cell fate decision seems normal in embryos with maternal *Ezh1/2* deletion.

The second cell lineage decision, occurring during implantation, segregates the ICM into EPI cells and primitive endoderm (PrE) cells that express Nanog and Gata6, respectively (Chazaud and Yamanaka, 2016). EPI cells produce the fetus, while PrE along with trophectoderm produces extraembryonic tissues (DePamphilis, 2016). The phenotypic similarity between dKO/+ embryos and *Nanog* null mice in peri-implantation (Mitsui et al., 2003) reminded us to examine whether loss of maternal *Ezh1/2* results in the dysregulation of second cell fate decision. As shown in Fig. 7, the morphology of E4.5 dKO/+ embryos had no discriminable difference compared to dF/+ embryos **(Fig. 7A)**. While the expression of Nanog and Gata6 were observed in late blastocysts **(Fig. 7B)**, Nanog-positive (Nanog+) cells in dKO/+ embryos were obviously fewer than that in dF/+ embryos, indicating EPI cells was reduced **(Fig. 7B)**. Cell counts showed that cell numbers of dF/+ and dKO/+ embryos were nearly equivalent **(Fig. 7C)**, while EPI cells in dKO/+ embryos were remarkably reduced (6.83 ± 3.24) compared to that in dF/+ embryos (11.58 ± 3.00) **(Fig. 7C)**. Unexpectedly, PrE cell number was significantly decreased, leaving about 66% to that of dF/+ embryos **(Fig. 7C)**. Moreover, both the ratio of EPI and PrE were obviously reduced **(Fig. 7D)**. TE cell number was not significant changed but its ratio to total cells was obviously increased, while mix-expressed cells appeared and accounted for about 4.21% of total cells **(Fig. 7C and 7D)**. Consistent with this result, Oct4 and Gata6 were expressed in late blastocysts **(sFig. 6A)**. Average cell numbers per embryo were not changed significantly **(sFig. 6B)**, but number of Oct4+ only cells were obviously decreased in dKO/+ embryos at E4.5 **(sFig. 6C and 6D)**. These results show that maternal *Ezh1/2* deletion impaired the second cell fate decision with less EPI cells and increased mix-expression of stem cell markers during implantation.

**Figure 7.**
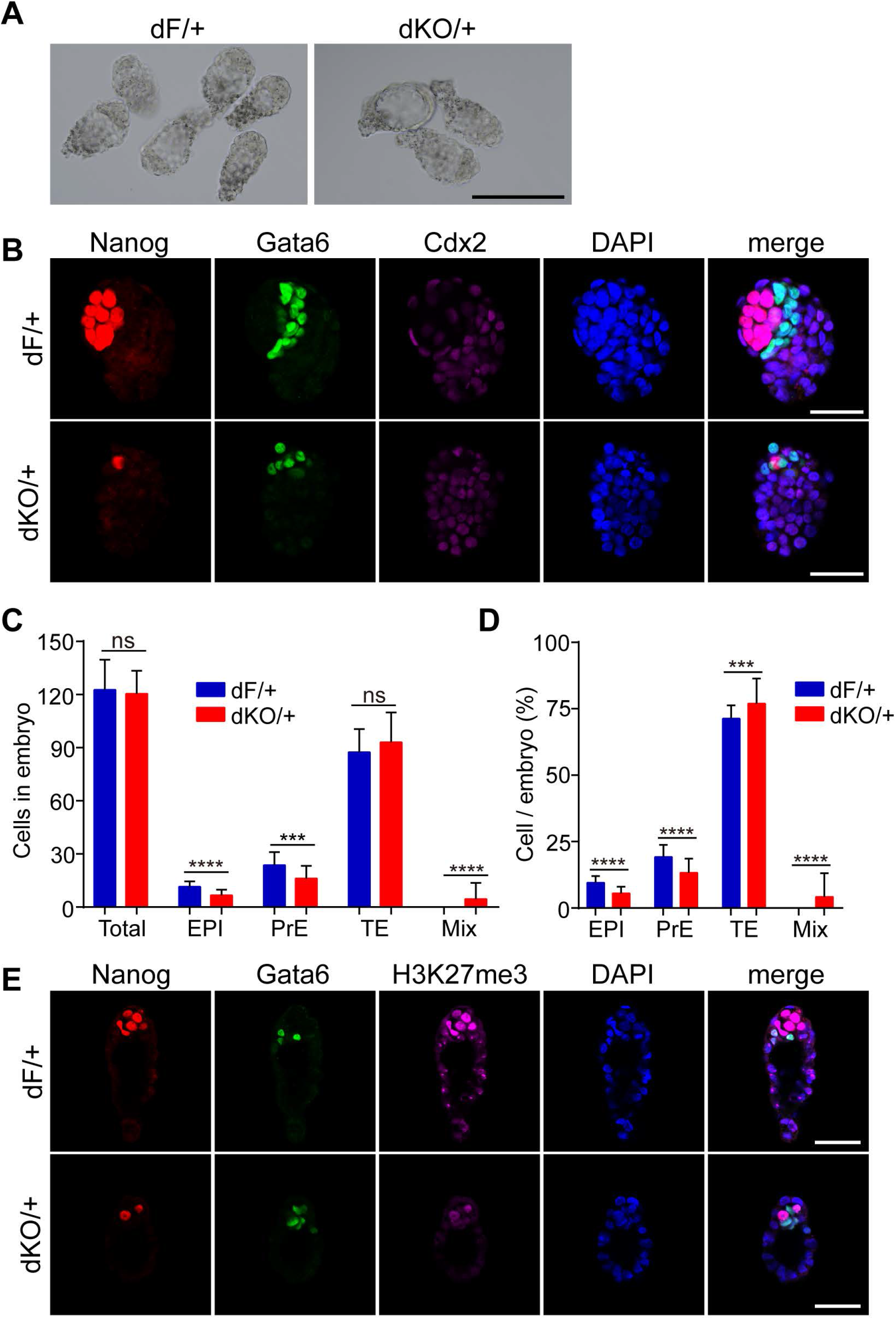
Decreased epiblast cells in dKO/+ Embryos at E4.5. **(A)** Representative images of flushed embryos at E4.5. Scale bars, 50 μm. **(B)** Representative images of stem cell distribution in late blastocysts at E4.5. EPI cells were stained with Nanog (red), PrE cells were stained with Gata6 (green) and TE cells were stained with Cdx2 (magenta). Number of total embryos: dF/+, n=31; dKO/+, n=35. Scale bars, 50 μm. **(C)** The number of different cell parts in embryos. Results are presented as mean ± SD. Statistical comparisons of values were made using Mann Whitney test (Total, EPI, PrE and Mix) or unpaired *t* test (TE). ns, not significant. \*\*\**P* < 0.001; \*\*\*\**P* < 0.0001. **(D)** The ratio of different cell parts. Results are presented as mean ± SD. Statistical comparisons of values were made using Mann Whitney test (PrE, TE and Mix) or unpaired *t* test (EPI). ns, not significant; \*\*\**P* < 0.001; \*\*\*\**P* < 0.0001. **(E)** Single optical sections of late blastocysts immunostained with Nanog, Gata6 and H3K27me3. Number of total embryos: dF/+, n=11; dKO/+, n=7.

It has been shown that the promoters of developmental genes are strongly marked with H3K27me3 in EPI in post-implantation embryos (Rugg-Gunn et al., 2010; Zheng et al., 2016), so we hypothesized that the defective second lineage commitment may be associated with reduced H3K27me3 signals. To test this hypothesis, we examined H3K27me3 modification and Nanog expression in embryos at E4.5. Besides H3K27me3 dot signals in some embryos, H3K27me3 modification appeared in Nanog+ cells and were reduced in dKO/+ embryos, without occurrence in Gata6+ cells of both dF/+ and dKO/+ late blastocysts **(Fig. 7E)**, indicating that EPI development is associated with H3K27me3 modification.

In summary, the maternal Ezh1/2 is crucial for second cell lineage determination and propagation of EPI state during implantation.

## Discussion

Our study investigated the effects of maternal *Ezh1/2* on H3K27 methylation patterns and embryonic development by deletion of *Ezh1* and *Ezh2* in oocytes in mice. The results showed that *Ezh2* is required for H3K27me3 and *Ezh1* partially complements *Ezh2* for H3K27me2 in oocytes. The effect of maternal *Ezh1/2* was inherited in descendant embryos. Loss of maternal *Ezh1/2* causes delayed restoration of H2K27me2/3 modification in preimplantation embryos *in vivo*. We also demonstrate that loss of maternal *Ezh1/2* has long-term developmental consequences resulting in growth retardation and sub-lethality in mice. The maternal *Ezh1/2* is essential for second lineage determination and EPI cell fate during implantation. These findings uncover the essential function of maternal *Ezh1/2* on H3K27me2/3 modification and embryonic development, which provides a novel understanding in embryonic development regulated by epigenetic modulators.

### Maternal *Ezh1* and *Ezh2* in descendant development in mice

It is shown that Ezh1 and Ezh2 are functionally redundant (Ezhkova et al., 2009; Margueron et al., 2008; Shen et al., 2008), and tissues deficient of *Ezh1* and *Ezh2* show severe defects than that lack of *Ezh2* (Ezhkova et al., 2011; Mochizuki-Kashio et al., 2015). This is the case in the context in our work. Mice lacking *Ezh1* were normal (Ezhkova et al., 2011), while oocyte-specific deletion of *Ezh2* induces growth retardation in offspring but does not influence mouse fertility (Erhardt, 2003), which is consistent with the fertility performance of dCtrl and sKO females in the present work. In our mice model, double knockout of *Ezh1* and *Ezh2*, but not *Ezh1* or *Ezh2* alone deficiency in oocytes caused embryonic sub-lethality, suggesting *Ezh1* complements (or partially) *Ezh2* function in embryonic development and maternal *Ezh2* combined with *Ezh1* governs the descendant development. Notably, we notice that patterns of H3K27me3 in late blastocysts and the breeding results are consistent to some extent. Considering that H3K27me3 modification is associated with pluripotent EPI, it is possible that the pattern of H3K27me3 in peri-implantation may determine the potential of embryonic development, which might explain fertility discrepancy between different groups.

### Maternal *Ezh1* and *Ezh2* in H3K27 methylation in pre-implantation embryos *in vivo*

Our knockout mice model show that Ezh2 is required for H3K27me3 and *Ezh1* complements *Ezh2* in H3K27me2 in oocytes. The effect of *Ezh1/2* is inherited in maternal knockout zygotes. H3K27me3 was lost in sKO/+ and dKO/+ maternal PN while H3K27me2 was reduced in sKO/+ female PN but absent in dKO/+ female PN. H3K27me2 signal was almost undetectable in both paternal and maternal PN in dKO/+ zygotes, which is similar to H3K27me2 pattern in Eed m-p+ zygotes, but different with a prior report in which H3K27me2 of maternal PN is normal in Ezh2m-p+ *Ezh1* siRNA zygotes (Meng et al., 2020). In addition, the asymmetric signal of H3K27me2/3 in parental PN of sF/+ and dF/+ zygotes are in coincidence with previous reports (Huang et al., 2014; Santos et al., 2005). Interestingly, similar patterns of H3K27me3 and H3K27me2 signal appear in paternal PN of maternal KO zygotes in this work. Normally, paternal PN is initially histone hypo-methylated and gradually deposited with H3K27me2/3 modifications in late zygotes, due to a preferential recruitment of PRC2 complex to maternal PN and apparent in paternal PN at later stage (Burton and Torres-Padilla, 2010; Erhardt, 2003). The lost deposition of H3K27me2/3 in paternal PN of KO/+ zygotes indicates PRC2 complex is inactive in late zygotes or at least absent in paternal PN. According to these results, we speculate that, in our study, *Ezh1* and *Ezh2* are required for *de novo* H3K27me2 in oocytes which could be restored and inherited in maternal PN of zygotes.

Deficiency of Ezh1 does not affect H3K27me2/3 patterns, while loss of maternal Ezh2 alone or combined with Ezh1 leads to delayed restoration of H3K27me2/3 in preimplantation embryos *in vivo*, indicating that maternal Ezh2-mediated H3K27me2/3 were inherited in early embryos. H3K27me2/3 status in dKO/+ embryos is severely delayed than that in sKO/+, suggesting Ezh1 compensating to sustain some residual PRC2 activity (Mochizuki-Kashio et al., 2015). However, H3K27me2/3 patterns in *in vivo* preimplantation embryos in our study are different with previous reports (Erhardt, 2003; Inoue et al., 2018; Meng et al., 2020). H3K27me3 was not recovered in early blastocysts of sKO/+ and dKO/+ embryos in our research. Erhardt et al. showed that emergence of normal H3K27me3 pattern in maternal Ezh2 knockout embryos (16-cell stage) is beyond embryonic Ezh2 activation (Erhardt, 2003), while H3K27me3 of maternal Eed knockout embryos becomes comparable at the morula stage (Inoue et al., 2018). Reasons for this difference may be as follows: (1) different collection of embryos, as histone modification may be changed by *in vitro* operation and culture (Kohda, 2013); (2) activation of Ezh1/2 may be later than Eed in maternal knockout embryos. Additionally, the conversation of histone methylation to H3K27me3 is more time consuming compared to H2K27me1 and H3K27me2 (Anne Laugesen et al., 2019), and a recent work indicates that H3K27me2 might be a critical prerequisite for de novo H3K27me3 (Meng et al., 2020). So, the more delayed restoration of H3K27me3 may be induced by deferred platform of H3K27me2. In dKO/+ preimplantation embryos, H3K27me2 appeared from morula stage and completely recovered until late blastocysts. Since Ehmt1 is also involved in the establishment of H3K27me2 (Meng et al., 2020), we suppose that this may be associated with delayed embryonic Ezh1/2 or Ehmt1 transcription for initiating H3K27me2. This hypothesis should be verified in the future work.

It’s worth noting that dKO/+ morulae had more H3K27me2 dot signals, which is reminiscent of the intense staining and the role of *Xist* and H3K27me3 in XCI (Borensztein et al., 2017; Chen and Zhang, 2020; Strehle and Guttman, 2020), indicating that H3K27me2 may act on XCI in early embryos. In addition, Inoue et al. showed that maternal Eed knockout causes active of maternal X chromosome (Xm) and decreased expression level of Xm-linked genes in morula, but the initial repression of X-linked genes might be independent of H3K27me3 (Inoue et al., 2018). It’s known that H3K27me2 is a repressive modification (Conway et al., 2015), so we guess H3K27me2 might be involved in the repression of Xm.

Above all, the capacity of H3K27 methylation reprogramming in preimplantation embryos was compromised by deletion of maternal Ezh2 alone or combined with Ezh1.

### Maternal PRC2-H3K27me3 in second cell fate decision

Erhardt et al. suggested that Ezh2 may be crucial for the propagation of the pluripotent state and cell fate determination during early stages of embryo (Erhardt, 2003). In this work, we reveal that maternal loss of Ezh1/2 leads to reduced EPI cells and defective second cell fate specification during implantation, unveiling the importance of Ezh1/2 and H3K27me3 for the EPI state and cell fate decision. Therefore, the fact that stem cells from Ezh2 mutant embryo are difficult be established and show impaired potential for outgrowth (O’Carroll et al., 2001) may associate with failed pluripotent EPI cells induced by abolishment of Ezh2 and H3K27me3.

It’s now clear that reduction of ICM cell numbers diminishes the potential of embryogenesis (Tam, 1988). The initiation of gastrulation is suggested to need attainment of either a threshold of cell number or tissue mass of EPI (Kojima et al., 2014). Our results showed markedly decreased ICM/EPI cells in dKO/+ embryos during implantation. It is likely that deficient ICM/EPI development could be responsible for embryonic loss and growth retardation during post-implantation. It is worth mentioning that the *in vitro* culture (IVC) system is a powerful tool for visualizing EPI development and its surrounding tissues through the implantation stages (Bedzhov and Zernicka-Goetz, 2014; Morris et al., 2012). By this system, the dynamics of blastocyst transforming into the egg cylinder will be directly revealed.

In our study, EPI cells (Nanog+) possess H3K27me3 signal in both dF/+ and dKO/+ embryos at E4.5. Similar H3K27me3 patterns are observed in previous reports (Erhardt, 2003; Liu et al., 2019). In ESCs, a set of genes are bivalent modified with H3K4me3 and H3K27me3, and stay in “poised” (Liu et al., 2016; Rugg-Gunn et al., 2010). Withdrawal of PRC2 core components and H3K27me3 methylation results in de-repression of bivalent genes and abnormal differentiation (Azuara et al., 2006; Boyer et al., 2006; Jorgensen et al., 2006; Lee et al., 2006). In consideration of H3K27me3 modifications on developmental genes in EPI cells (Rugg-Gunn et al., 2010; Zheng et al., 2016), decreased Nanog+ cells may be resulted from the deficiency of H3K27me3 and the de-repression of cell fate specific factors, which lead to differentiation into other lineages or mix-expression. A recent report demonstrates that in mouse ESCs, Nanog blocks differentiation by maintaining H3K27me3 modification at developmental regulators (Heurtier et al., 2019). However, whether this regulatory network functions *in vivo* remains to be elaborately addressed. For the two rounds of cell fate specification, cells in the wrong position or with ectopic expression of transcriptional profile will eventually be cleared out by apoptosis (Zhu and Zernicka-Goetz, 2020). Whether the decreased number of ICM (EPI and PrE) cells is associated with apoptosis needs further assessment.

In brief, loss of maternal Ezh1/2 and delayed restoration of H3K27 methylation change the epigenetic plasticity of pluripotent cells.

### Maternal Ezh1/2 in placental development

It’s known that lack of chorioallantoic attachment and branching results in placental failure and embryonic mortality (Rossant and Cross, 2001; Walentin et al., 2016). Hence, in our work, failure of placentation in mid-gastrulation is at least one of the reasons for embryonic loss. But it’s not clear how loss of maternal Ezh1/2 leads to failed chorioallantoic attachment, or it might be a by-product by additional mechanisms (Inoue et al., 2018; Perez-Garcia et al., 2018), which needs further investigation. In contrast, enlarged placentas existed in late development of dKO/+ embryos, which was characterized by expansion of spongiotrophoblast layer. This phenotype was reported in mouse embryos of somatic cell nuclear transfer (SCNT) (Tanaka et al., 2001). Some recent reports have demonstrated that loss of H3K27me3 imprinting in SCNT contributes to placentomegaly of SCNT embryos, and placenta weight could be ameliorated by regulating H3K27me3-imprinted genes (Inoue et al., 2020; Matoba et al., 2018; Wang et al., 2020). Given that maternal Eed knockout causes loss of H3K27me3 imprinting in the extraembryonic genes (Inoue et al., 2018), it’s possible that loss of maternal Ezh1/2 may contribute to the loss of H3K27me3 dependent imprinting, thus the enlarged placenta is probably caused by derepression of H3K27me3-imprinted genes.

## Materials and Methods

### Animals

Unless otherwise stated, mice used in this study were of mixed background (129/SvJ and C57BL/6). The *Ezh2 loxp* mice were gifts from Professor Alexander Tarakhovsky (The Rockefeller University, New York, USA). Ezh1 knockout mice were obtained from Thomas Jenuwein’s laboratory and will be reported elsewhere. Mice were housed at Xiamen University laboratory Animal Center using a 12 h light-dark cycle. Food and water were available *ad libitum* and room temperature was 24 °C with controlled humidity. All animal work was undertaken in accordance with Xiamen University Animal Ethics Committee approvals (2015014). Genotyping was performed from tail DNA. Primers for genotype were in **Supplement Table S2**.

### Fertility studies

Fertility studies were performed as previously described (Andreu-Vieyra et al., 2008). To evaluate reproductive performance, individually caged females (2 months old) were bred with adult wild type males (C57BL/6J) of known fertility. The number of litters and the number of pups were recorded over a 6-month period.

### Mating, implantation, embryo collection and decidua dissection

When basic embryological methods performed, adult dCtrl and dKO females were naturally mated with adult wild type males (C57/BL6J). The morning of finding plug was counted as embryonic day 0.5 (E0.5) or day 1 of pregnancy. Implantation sites on day 5 (09:00 am) of pregnancy were visualized by an intravenous injection of □hicago blue dye solution. Blastocysts were flushed from uterine horn with M2 medium at E3.75 and E4.5. Deciduae of pregnant females were dissected from E4.75-E10.5. Fetus and placentas were dissected carefully under stereoscope at E10.5 and E17.5.

### Histology, immunohistochemistry and immunofluorescence

Tissues were fixed in 4% paraformaldehyde (PFA), dehydrated in ethanol, embedded in paraffin wax, and sectioned (5 μm). To observe the morphology of tissues, sections were stained with the hematoxylin-eosin (HE). For immunohistochemistry, sections were treated with 3% H_2_O_2_ for 20 minutes after antigen retrieval, followed by 1 hour in 5% bovine serum albumin (BSA). Sections incubated in primary antibodies overnight at 4 °C were subsequently treated with PV-9001 kits (ZSBIO) before being exposed to diaminobenzidine (DAB). Finally, sections were counterstained with hematoxylin. For immunofluorescence, after antigen retrieval, sections were blocked in 5% bovine serum albumin or 5% donkey serum and then incubated in primary antibodies overnight at 4 °C. Then the sections were washed in wash buffer and incubated with the appropriate Alexa-Fluor-conjugated secondary antibodies at 37 °C for 1 h. After staining with DAPI (Vector laboratories), samples were analyzed using confocal microscopy (Zeiss LSM 780). Antibodies were in **Supplement Table S3**.

### Oocytes and early embryos collection

MII oocytes were collected from females (3-5 weeks old) super-ovulated by injection with PMSG and hCG (San-Sheng pharmaceutical Co. Ltd). Preimplantation embryos were collected from super-ovulated females mated with adult wild type males (C57BL/6). Each set of embryos at a particular stage was flushed from the reproductive tract at defined time periods after hCG administration: 20 h (MII oocyte), 27-28 h (late zygote), 48 h (late 2-cell), 54-56 h (4-cell), 68-70 h (8-cell), 80 h (morula), 90-96 h (early blastocyst) and 120 h (late blastocyst).

### Whole-mount immunostaining

Oocytes and embryos were fixed in 4% PFA containing 0.1% BSA for 20 min at room temperature. After that, all fixed samples were washed by PBS containing 0.1% BSA and permeabilized in 0.1% Triton X-100 in PBS with 0.1% BSA at room temperature for 30 min, and blocked in 1% BSA or 5% donkey serum in PBS at room temperature for 1 h. Blocked oocytes and embryos were incubated with primary antibodies at 4 °C overnight. Secondary antibodies were incubated at 37 °C for 1 h. After washing, they were mounted on a glass slide with DAPI. Fluorescence was detected under a laser-scanning confocal microscope (Zeiss, LSM 780).

### Antibody-labeling

4AF647R-Antibody conjugation Kit (4A Biotech) was used to label H3K27me3 antibody with 4AF647R Dyes. According to the labeling protocol, H3K27me3 antibody solutions (CST, 9733) were concentrated by ultrafiltration and were resuspended in PBS to 0.5-1 mg/ml. An appropriate amount of antibody to be labeled was transferred to a clean tube. 1/10 volume of Reaction Buffer was added to the tube with antibody and then mixed well. The vial was incubated in the dark for 30 min at room temperature. Labeled antibody solution was diluted with Storage Buffer, and stored in single use aliquots at -20 °C. When performing immunofluorescence, labeled H3K27me3 was used after standard immunofluorescence staining with primary and secondary antibodies.

### Quantitative real time PCR (QPCR) analyses

Oocyte preparation was as reported (Kim et al., 2015). RNA from oocytes was isolated using the SuperPrep Cell Lysis & RT Kit for real-time PCR (TOYOBO, Osaka, Japan). PCR was performed on the Agilent AriaMx Real-Time PCR System. Primers were in **Supplement Table S2**. The relative number of transcripts was calculated by the cycle threshold method as described by Agilent using the AtiaMx Real-Time PCR System Software and normalized to the endogenous reference (*β*-*actin*). The relative amount of target gene expression for each sample was calculated and plotted as the mean ± SD.

### Western blot analyses

Western blots were carried out as previously described (Kim et al., 2015). Equal oocytes were collected in PBS containing 0.1% BSA. Proteins were isolated by centrifuging oocytes at 12,000 g for 15 minutes at 4 °C. Protein samples were then mixed and treated with 2×SDS buffer containing protease inhibitor. Protein molecules were separated by gel electrophoresis and transferred onto PVDF membranes. Membranes were blocked in 5% milk in TBST and incubated in primary antibodies. Membranes were incubated in second antibodies after washing. Band was detected by using Bio-Rad imaging systems.

### Vasectomy

Vasectomy was operated as previous documented (Bermejo-Alvarez et al., 2014). Wild type male mice (CD1) with a proven mating performance were used and anesthetized. Abdomen surfaces were cleaned and wiped with 70% ethanol. Abdomen was opened and vas deference was exposed by grabbing the testicular adipose pad with forceps. The vas deferens was cut and cauterize in two points at once by using flame dressing forceps. Testicle, epididymis and vas deferens were moved back to the abdominal cavity. The other vas deference was proceeded. Muscle and skin were sutured. The vasectomized male was moved to the cage placed on a warm stage and observed until it recovered from anesthesia. Wound clips were removed 14 days after vasectomy. The infertility of the vasectomized male was tested by mating fertile females before using it to obtain recipients.

### Embryo transfer

Embryo transfer was performed as previously described (Bermejo-Alvarez et al., 2014). Female mice were super-ovulated and mated with wild type males (C57BL/6). Blastocysts were flushed out of the uteri on day 4 of pregnancy, then transferred into the uteri of pseudo-pregnant female mice (CD1, wild type) mated with vasectomized males. About 17 days after embryo transfer, the recipients were examined for the presence of fetus. The number of fetus was recorded.

### Statistical analyses and data visualization

Statistical analyses were performed in GraphPad Prism 7. Quantitative data presented, shows the mean ± SD, percentages, or the total number of data points obtained. Levels of significance were calculated using Mann Whitney test, unpaired *t*-test or *χ*^2^ test. In all figures: ns, not significant; *, *P* < 0.05; **, *P* < 0.01; ***, *P* < 0.001; ****, *P* < 0.0001.

## Competing of interest

The authors declare no conflict of interest.

## Funding

This work was supported by the National Key Research and Development Program of China (grant no. 2018YFC1003701 and 2017YFC1001402) and the National Natural Science Foundation of China (grant no. 31970797 and 31671564).

## Data availability

All data generated or analyzed during this study are included in this published article and its supplementary information files

## Author contributions

Z. Lu and H. Wang conceived the original ideas, designed the project, and wrote the manuscript with inputs from Y. Zhao and S. Gao. Y. Zhao performed the majority of the experiments with participation from D. Zhang, M. Liu, J. Lu and Y. Tian. All authors read and approved the final manuscript.

## Notes

### Competing Interest Statement

The authors have declared no competing interest.

## Reference

Andreu-Vieyra, C., Chen, R., Matzuk, M.M., 2008. Conditional deletion of the retinoblastoma (Rb) gene in ovarian granulosa cells leads to premature ovarian failure. Molecular endocrinology 22, 2141–2161.

Anne Laugesen, Jonas Westergaard Højfeldt, Helin, a.K., 2019. Molecular Mechanisms Directing PRC2 Recruitment and H3K27 Methylation. Molecular cell 74, 8–18.

Azuara, V., Perry, P., Sauer, S., Spivakov, M., Jorgensen, H.F., John, R.M., Gouti, M., Casanova, M., Warnes, G., Merkenschlager, M., Fisher, A.G., 2006. Chromatin signatures of pluripotent cell lines. Nat Cell Biol 8, 532–538.

Bedzhov, I., Zernicka-Goetz, M., 2014. Self-organizing properties of mouse pluripotent cells initiate morphogenesis upon implantation. Cell 156, 1032–1044.

Bermejo-Alvarez, P., Park, K.E., Telugu, B.P., 2014. Utero-tubal embryo transfer and vasectomy in the mouse model. J Vis Exp, e51214.

Borensztein, M., Okamoto, I., Syx, L., Guilbaud, G., Picard, C., Ancelin, K., Galupa, R., Diabangouaya, P., Servant, N., Barillot, E., Surani, A., Saitou, M., Chen, C.J., Anastassiadis, K., Heard, E., 2017. Contribution of epigenetic landscapes and transcription factors to X-chromosome reactivation in the inner cell mass. Nat Commun 8, 1297.

Boyer, L.A., Plath, K., Zeitlinger, J., Brambrink, T., Medeiros, L.A., Lee, T.I., Levine, S.S., Wernig, M., Tajonar, A., Ray, M.K., Bell, G.W., Otte, A.P., Vidal, M., Gifford, D.K., Young, R.A., Jaenisch, R., 2006. Polycomb complexes repress developmental regulators in murine embryonic stem cells. Nature 441, 349–353.

Bracken, A.P., Dietrich, N., Pasini, D., Hansen, K.H., Helin, K., 2006. Genome-wide mapping of Polycomb target genes unravels their roles in cell fate transitions. Genes & development 20, 1123–1136.

Burton, A., Torres-Padilla, M.E., 2010. Epigenetic reprogramming and development: a unique heterochromatin organization in the preimplantation mouse embryo. Brief Funct Genomics 9, 444–454.

Canovas, S., Ross, P.J., 2016. Epigenetics in preimplantation mammalian development. Theriogenology 86, 69–79.

Cao, R., Wang, L., Wang, H., Xia, L., Erdjument-Bromage, H., Tempst, P., Jones, R.S., Zhang, Y., 2002. Role of histone H3 lysine 27 methylation in Polycomb-group silencing. Science 298, 1039–1043.

Chazaud, C., Yamanaka, Y., 2016. Lineage specification in the mouse preimplantation embryo. Development 143, 1063–1074.

Chen, Z., Zhang, Y., 2020. Maternal H3K27me3-dependent autosomal and X chromosome imprinting. Nat Rev Genet 21, 555–571.

Conway, E., Healy, E., Bracken, A.P., 2015. PRC2 mediated H3K27 methylations in cellular identity and cancer. Curr Opin Cell Biol 37, 42–48.

DePamphilis, M.L., 2016. Preface, Mammalian Preimplantation Development, pp. xiii–xxi.

Erhardt, S., 2003. Consequences of the depletion of zygotic and embryonic enhancer of zeste 2 during preimplantation mouse development. Development 130, 4235–4248.

Ezhkova, E., Lien, W.H., Stokes, N., Pasolli, H.A., Silva, J.M., Fuchs, E., 2011. EZH1 and EZH2 cogovern histone H3K27 trimethylation and are essential for hair follicle homeostasis and wound repair. Genes & development 25, 485–498.

Ezhkova, E., Pasolli, H.A., Parker, J.S., Stokes, N., Su, I.H., Hannon, G., Tarakhovsky, A., Fuchs, E., 2009. Ezh2 orchestrates gene expression for the stepwise differentiation of tissue-specific stem cells. Cell 136, 1122–1135.

Ferrari, K.J., Scelfo, A., Jammula, S., Cuomo, A., Barozzi, I., Stutzer, A., Fischle, W., Bonaldi, T., Pasini, D., 2014. Polycomb-dependent H3K27me1 and H3K27me2 regulate active transcription and enhancer fidelity. Molecular cell 53, 49–62.

Heurtier, V., Owens, N., Gonzalez, I., Mueller, F., Proux, C., Mornico, D., Clerc, P., Dubois, A., Navarro, P., 2019. The molecular logic of Nanog-induced self-renewal in mouse embryonic stem cells. Nat Commun 10, 1109.

Huang, X.J., Wang, X., Ma, X., Sun, S.C., Zhou, X., Zhu, C., Liu, H., 2014. EZH2 is essential for development of mouse preimplantation embryos. Reproduction, fertility, and development 26, 1166–1175.

Inoue, A., Chen, Z., Yin, Q., Zhang, Y., 2018. Maternal Eed knockout causes loss of H3K27me3 imprinting and random X inactivation in the extraembryonic cells. Genes & development 32, 1525–1536.

Inoue, K., Ogonuki, N., Kamimura, S., Inoue, H., Matoba, S., Hirose, M., Honda, A., Miura, K., Hada, M., Hasegawa, A., Watanabe, N., Dodo, Y., Mochida, K., Ogura, A., 2020. Loss of H3K27me3 imprinting in the Sfmbt2 miRNA cluster causes enlargement of cloned mouse placentas. Nat Commun 11, 2150.

Jambhekar, A., Dhall, A., Shi, Y., 2019. Roles and regulation of histone methylation in animal development. Nature reviews. Molecular cell biology 20, 625–641.

Jorgensen, H.F., Giadrossi, S., Casanova, M., Endoh, M., Koseki, H., Brockdorff, N., Fisher, A.G., 2006. Stem cells primed for action: polycomb repressive complexes restrain the expression of lineage-specific regulators in embryonic stem cells. Cell Cycle 5, 1411–1414.

Kim, J., Singh, A.K., Takata, Y., Lin, K., Shen, J., Lu, Y., Kerenyi, M.A., Orkin, S.H., Chen, T., 2015. LSD1 is essential for oocyte meiotic progression by regulating CDC25B expression in mice. Nat Commun 6, 10116.

Kohda, T., 2013. Effects of embryonic manipulation and epigenetics. J Hum Genet 58, 416–420.

Kojima, Y., Tam, O.H., Tam, P.P., 2014. Timing of developmental events in the early mouse embryo. Semin Cell Dev Biol 34, 65–75.

Lee, T.I., Jenner, R.G., Boyer, L.A., Guenther, M.G., Levine, S.S., Kumar, R.M., Chevalier, B., Johnstone, S.E., Cole, M.F., Isono, K., Koseki, H., Fuchikami, T., Abe, K., Murray, H.L., Zucker, J.P., Yuan, B., Bell, G.W., Herbolsheimer, E., Hannett, N.M., Sun, K., Odom, D.T., Otte, A.P., Volkert, T.L., Bartel, D.P., Melton, D.A., Gifford, D.K., Jaenisch, R., Young, R.A., 2006. Control of developmental regulators by Polycomb in human embryonic stem cells. Cell 125, 301–313.

Liu, J., An, L., Wang, J., Liu, Z., Dai, Y., Liu, Y., Yang, L., Du, F., 2019. Dynamic patterns of H3K4me3, H3K27me3, and Nanog during rabbit embryo development. Am J Transl Res 11, 430–441.

Liu, X., Wang, C., Liu, W., Li, J., Li, C., Kou, X., Chen, J., Zhao, Y., Gao, H., Wang, H., Zhang, Y., Gao, Y., Gao, S., 2016. Distinct features of H3K4me3 and H3K27me3 chromatin domains in pre-implantation embryos. Nature 537, 558–562.

Margueron, R., Li, G., Sarma, K., Blais, A., Zavadil, J., Woodcock, C.L., Dynlacht, B.D., Reinberg, D., 2008. Ezh1 and Ezh2 maintain repressive chromatin through different mechanisms. Molecular cell 32, 503–518.

Margueron, R., Reinberg, D., 2011. The Polycomb complex PRC2 and its mark in life. Nature 469, 343–349.

Matoba, S., Wang, H., Jiang, L., Lu, F., Iwabuchi, K.A., Wu, X., Inoue, K., Yang, L., Press, W., Lee, J.T., Ogura, A., Shen, L., Zhang, Y., 2018. Loss of H3K27me3 Imprinting in Somatic Cell Nuclear Transfer Embryos Disrupts Post-Implantation Development. Cell Stem Cell 23, 343–354.

Matsui, Y., Mochizuki, K., 2014. A current view of the epigenome in mouse primordial germ cells. Mol Reprod Dev 81, 160–170.

Meng, T.G., Zhou, Q., Ma, X.S., Liu, X.Y., Meng, Q.R., Huang, X.J., Liu, H.L., Lei, W.L., Zhao, Z.H., Ouyang, Y.C., Hou, Y., Schatten, H., Ou, X.H., Wang, Z.B., Gao, S.R., Sun, Q.Y., 2020. PRC2 and EHMT1 regulate H3K27me2 and H3K27me3 establishment across the zygote genome. Nat Commun 11, 6354.

Michalak, E.M., Burr, M.L., Bannister, A.J., Dawson, M.A., 2019. The roles of DNA, RNA and histone methylation in ageing and cancer. Nature reviews. Molecular cell biology 20, 573–589.

Mitsui, K., Tokuzawa, Y., Itoh, H., Segawa, K., Murakami, M., Takahashi, K., Maruyama, M., Maeda, M., Yamanaka, S., 2003. The homeoprotein Nanog is required for maintenance of pluripotency in mouse epiblast and ES cells. Cell 113, 631–642.

Mochizuki-Kashio, M., Aoyama, K., Sashida, G., Oshima, M., Tomioka, T., Muto, T., Wang, C., Iwama, A., 2015. Ezh2 loss in hematopoietic stem cells predisposes mice to develop heterogeneous malignancies in an Ezh1-dependent manner. Blood 126, 1172–1183.

Mole, M.A., Weberling, A., Zernicka-Goetz, M., 2020. Comparative analysis of human and mouse development: From zygote to pre-gastrulation. Curr Top Dev Biol 136, 113–138.

Morris, S.A., Grewal, S., Barrios, F., Patankar, S.N., Strauss, B., Buttery, L., Alexander, M., Shakesheff, K.M., Zernicka-Goetz, M., 2012. Dynamics of anterior-posterior axis formation in the developing mouse embryo. Nat Commun 3, 673.

O’Carroll, D., Erhardt, S., Pagani, M., Barton, S.C., Surani, M.A., Jenuwein, T., 2001. The polycomb-group gene Ezh2 is required for early mouse development. Molecular and cellular biology 21, 4330–4336.

Perez-Garcia, V., Fineberg, E., Wilson, R., Murray, A., Mazzeo, C.I., Tudor, C., Sienerth, A., White, J.K., Tuck, E., Ryder, E.J., Gleeson, D., Siragher, E., Wardle-Jones, H., Staudt, N., Wali, N., Collins, J., Geyer, S., Busch-Nentwich, E.M., Galli, A., Smith, J.C., Robertson, E., Adams, D.J., Weninger, W.J., Mohun, T., Hemberger, M., 2018. Placentation defects are highly prevalent in embryonic lethal mouse mutants. Nature 555, 463–468.

Prokopuk, L., Stringer, J.M., White, C.R., Vossen, R., White, S.J., Cohen, A.S.A., Gibson, W.T., Western, P.S., 2018. Loss of maternal EED results in postnatal overgrowth. Clin Epigenetics 10, 95.

Rossant, J., Cross, J.C., 2001. Placental development: lessons from mouse mutants. Nat Rev Genet 2, 538–548.

Rugg-Gunn, P.J., Cox, B.J., Ralston, A., Rossant, J., 2010. Distinct histone modifications in stem cell lines and tissue lineages from the early mouse embryo. Proceedings of the National Academy of Sciences of the United States of America 107, 10783–10790.

Saitou, M., Kagiwada, S., Kurimoto, K., 2012. Epigenetic reprogramming in mouse pre-implantation development and primordial germ cells. Development 139, 15–31.

Santos, F., Peters, A.H., Otte, A.P., Reik, W., Dean, W., 2005. Dynamic chromatin modifications characterise the first cell cycle in mouse embryos. Developmental biology 280, 225–236.

Schuettengruber, B., Cavalli, G., 2009. Recruitment of polycomb group complexes and their role in the dynamic regulation of cell fate choice. Development 136, 3531–3542.

Shen, X., Liu, Y., Hsu, Y.J., Fujiwara, Y., Kim, J., Mao, X., Yuan, G.C., Orkin, S.H., 2008. EZH1 mediates methylation on histone H3 lysine 27 and complements EZH2 in maintaining stem cell identity and executing pluripotency. Molecular cell 32, 491–502.

Simon, J.A., Kingston, R.E., 2009. Mechanisms of polycomb gene silencing: knowns and unknowns. Nature reviews. Molecular cell biology 10, 697–708.

Strehle, M., Guttman, M., 2020. Xist drives spatial compartmentalization of DNA and protein to orchestrate initiation and maintenance of X inactivation. Curr Opin Cell Biol 64, 139–147.

Tam, P.P., 1988. Postimplantation development of mitomycin C-treated mouse blastocysts. Teratology 37, 205–212.

Tam, P.P., Loebel, D.A., 2007. Gene function in mouse embryogenesis: get set for gastrulation. Nat Rev Genet 8, 368–381.

Tanaka, S., Oda, M., Toyoshima, Y., Wakayama, T., Tanaka, M., Yoshida, N., Hattori, N., Ohgane, J., Yanagimachi, R., Shiota, K., 2001. Placentomegaly in cloned mouse concepti caused by expansion of the spongiotrophoblast layer. Biology of reproduction 65, 1813–1821.

Walentin, K., Hinze, C., Schmidt-Ott, K.M., 2016. The basal chorionic trophoblast cell layer: An emerging coordinator of placenta development. Bioessays 38, 254–265.

Wang, L.Y., Li, Z.K., Wang, L.B., Liu, C., Sun, X.H., Feng, G.H., Wang, J.Q., Li, Y.F., Qiao, L.Y., Nie, H., Jiang, L.Y., Sun, H., Xie, Y.L., Ma, S.N., Wan, H.F., Lu, F.L., Li, W., Zhou, Q., 2020. Overcoming Intrinsic H3K27me3 Imprinting Barriers Improves Post-implantation Development after Somatic Cell Nuclear Transfer. Cell Stem Cell 27, 315–325.

Yan, J., Dutta, B., Hee, Y.T., Chng, W.J., 2019. Towards understanding of PRC2 binding to RNA. RNA Biol 16, 176–184.

Yu, J.R., Lee, C.H., Oksuz, O., Stafford, J.M., Reinberg, D., 2019. PRC2 is high maintenance. Genes & development 33, 903–935.

Zenk, F., Loeser, E., Schiavo, R., Kilpert, F., Bogdanović, O., Iovino, N., 2017. Germ line-inherited H3K27me3 restricts enhancer function during maternal-to-zygotic transition. Science 357, 212–216.

Zhang, B., Zheng, H., Huang, B., Li, W., Xiang, Y., Peng, X., Ming, J., Wu, X., Zhang, Y., Xu, Q., Liu, W., Kou, X., Zhao, Y., He, W., Li, C., Chen, B., Li, Y., Wang, Q., Ma, J., Yin, Q., Kee, K., Meng, A., Gao, S., Xu, F., Na, J., Xie, W., 2016. Allelic reprogramming of the histone modification H3K4me3 in early mammalian development. Nature 537, 553–557.

Zheng, H., Huang, B., Zhang, B., Xiang, Y., Du, Z., Xu, Q., Li, Y., Wang, Q., Ma, J., Peng, X., Xu, F., Xie, W., 2016. Resetting Epigenetic Memory by Reprogramming of Histone Modifications in Mammals. Molecular cell 63, 1066–1079.

Zhu, M., Zernicka-Goetz, M., 2020. Principles of Self-Organization of the Mammalian Embryo. Cell 183, 1467–1478.

